# Replication of Equine arteritis virus is efficiently suppressed by purine and pyrimidine biosynthesis inhibitors

**DOI:** 10.1101/2020.04.10.035402

**Authors:** José Carlos Valle-Casuso, Delphine Gaudaire, Lydie Martin-Faivre, Anthony Madeline, Patrick Dallemagne, Stéphane Pronost, Hélène Munier-Lehmann, Stephan Zientara, Pierre-Olivier Vidalain, Aymeric Hans

## Abstract

RNA viruses are responsible for a large variety of animal infections. Equine Arteritis Virus (EAV) is a positive single-stranded RNA virus member of the family *Arteriviridae* from the order *Nidovirales* like the *Coronaviridae.* EAV causes respiratory and reproductive diseases in equids. Although two vaccines are available, the vaccination coverage of the equine population is largely insufficient to prevent new EAV outbreaks around the world. In this study, we present a high-throughput *in vitro* assay suitable for testing candidate antiviral molecules on equine dermal cells infected by EAV. Using this assay, we identified three molecules that impair EAV infection in equine cells: the broad-spectrum antiviral and nucleoside analog ribavirin, and two compounds previously described as inhibitors of dihydroorotate dehydrogenase (DHODH), the fourth enzyme of the pyrimidine biosynthesis pathway. These molecules effectively suppressed cytopathic effects associated to EAV infection, and strongly inhibited viral replication and production of infectious particles. Since ribavirin is already approved in human and small animal, and that several DHODH inhibitors are in advanced clinical trials, our results open new perspectives for the management of EAV outbreaks.

## 1. Introduction

Viruses with a RNA genome infect both animals and plants, and dominate the eukaryotic virome by their diversity and deleterious effects ^1^. Indeed, RNA viral infections are a substantial public health burden affecting humans and domestic animals ^2^. The pandemic of SARS-CoV-2 coronavirus is the latest and most striking example with thousands of deaths reported so far in different countries in Europe, Asia and Middle East. Horses are not an exception, and five of seven equine viruses listed by the World Organization for Animal Health are RNA viruses (OIE). Unfortunately, the veterinary therapeutic arsenal to fight against equine viral diseases is limited, or even not-existent as no antiviral molecule is listed today. During the last years, several studies have used high-throughput screening (HTS) techniques to scan small compound libraries that could impair the replication of equine viruses. These studies were mostly focused on zoonotic viruses like West-Nile virus ^3,4^. Non-zoonotic equine RNA viruses, such as equine arteritis virus (EAV), were not included in those screenings. Equine arteritis virus (EAV) is a positive-sense single-stranded RNA virus, member of the subfamily *Equarterivirinae*, member of the family *Arteriviridae*, genus *Alphaarterivirus* from order *Nidovirales* like the *Coronaviridae*. Equine viral arteritis disease is a recurrent viral infection that causes important economic losses to the horse industry. In most of the cases, EAV infection is subclinical, but some viral strains are virulent and induce fever with body temperature going up to 41 °C, depression, anorexia, edema, nasal discharge, and conjunctivitis. EAV is a virus transmitted by respiratory or venereal routes. After EAV primary infection, up to 70% of stallions may become persistently infected and shed the virus in their semen. Those stallions, also called shedder stallions, are a key component in EAV epidemiology and transmission of the disease ^5,6^. Moreover, infected pregnant mares could abort or stillbirth a weak foal. Besides, foal infection can cause fulminating pneumonia with a lethal outcome. The loss of these animals, together with the restrictive reproductive activity of infected stallions, are significant complications for the equine industry ^7^.

Previous works exploring EAV inhibitors have proposed to use morpholino or peptide-conjugated morpholino oligomers specifically designed against the EAV genome as a therapy to impair viral replication ^8^. However, this promising therapy is not easily applicable for use *in vivo* in large animal such as horses. Although, two effective EAV vaccines have been available since 1985, the proportion of horses vaccinated against EAV in the equine population remains insufficient to control the disease ^9^. We think that a complementary approach using antiviral molecules would be ideal to complement vaccination, reduce viral spreading in equine structures and contain EAV outbreaks. This is supported by a recent study showing that a global strategy based on vaccination combined with drug therapies enhances protection against foot-and-mouth disease ^10^. In our study, we have developed a high-throughput cell-viability assay based on infected equine dermal cells. Then, we took advantage of this new assay to explore the potential of broad-spectrum antivirals (BSA) against EAV infection. BSA are small molecules that, because of their peculiar mode of action, impair the replication of multiple viruses from different species and genotypes. Nucleotide and nucleoside analogs are excellent examples of BSA. They inhibit transcription and/or replication of different RNA and DNA viruses by targeting components of the viral replication machinery. However, it is established that numerous viruses eventually mutate and develop resistance to these molecules ^11-13^. To circumvent this limitation, newly designed BSA overtake the drug resistance problem by targeting host cellular pathways hijacked by the virus to replicate instead of viral factors themselves.

Interestingly, one of the most studied BSA is ribavirin which has been described as an inhibitor of porcine reproductive and respiratory syndrome virus (PRRSV) and porcine epidemic diarrhea virus (PEDV) ^14^, two nidoviruses closely related to EAV. *In vivo*, ribavirin is used for treating humans or animals infected with emerging viruses such as Lassa virus or Hantavirus for which therapeutic options are limited ^15-18 15,19-21^. Ribavirin is a synthetic analog of guanosine that interferes with viral replication through multiple direct and indirect mechanisms. In particular, ribavirin inhibits inosine 5’-monophosphate dehydrogenase (IMPDH), a key enzyme catalyzing the first committed and rate-limiting step in the *de novo* synthesis of guanine nucleotides. As such, IMPDH inhibition interferes with the production of GTP that is necessary for the synthesis of viral RNA molecules ^22^, but other mechanisms have been reported such as the direct inhibition of viral polymerases or the lethal mutagenesis of viral genomes ^23,24^. Besides ribavirin, we also tested two newly designed BSA that target the pyrimidine biosynthesis pathway, namely GAC50 and IPPA17-A04. These two molecules are inhibitors of dihydroorotate dehydrogenase (DHODH), the fourth and rate-limiting enzyme of the *de novo* pyrimidine biosynthesis pathway, and were shown to impair the replication of many different positive-strand and negative-strand RNA viruses ^25-27^. In this study, we have shown, for the first time, that both ribavirin and pyrimidine biosynthesis inhibitors inhibit EAV in equine dermal cell cultures.

## 2. Results

### 2.1. Development of a microplate assay for monitoring EAV-induced cytopathic effects

EAV is a cytopathic virus. In this study, we took advantage of this specificity to develop a microplate assay for the identification of EAV inhibitors. As a cellular model, we selected the ED cell line that originates from equine dermis and thus matches the host specificity of EAV. To setup our *in vitro* assay, we first analyzed the proliferation of non-infected ED cells in 96-well plates. To determine the number of viable cells in each culture well, we quantified ATP which parallels the number of metabolically active, living cells. Assays were performed using a commercial reagent based on the principle that *firefly* luciferase luminescence is proportional and dependent on ATP concentrations (CellTiter-Glo Luminescent Cell Viability Assay; Promega). In non-infected cells, ATP levels increased by 41% in the first 24 h as a consequence of cellular proliferation, but then plateaued at 48 h as cell cultures reached confluence (Figure 1A). Same experiment was performed using cell cultures infected with the Bucyrus strain of EAV at a multiplicity of infection (MOI) of 0.5. As shown in Figure 1A, ATP levels in infected cultures only increased by 22% during the first 24 h as a consequence of infection, and then collapsed after 48 h of culture. When cells were analyzed by phase-contrast microscopy, infected and non-infected cultures looked similar at 24 h, but most cells in infected wells were dead at 48 h post-infection (h.p.i) as a result of EAV cytopathic effects (data not shown). We also confirm that observed cytopathic effects reflect EAV replication by quantifying the number of viral genome copies at 1, 24 h and 48 h post-infection using RT-qPCR. Results confirmed EAV viral replication in ED cells as the number of genome copies increased by more than 2 logs in the first 24 h, and by 3 logs at 48 h post-infection (Figure 1B). Finally, we showed that infectious viral particles are also produced by ED cells. Indeed, at least 1×10^5^ PFU/ml were detected in supernatants of infected cultures 48 h.p.i. (Figure 1C). Therefore, ED cells represent an excellent *in vitro* model for EAV infection, and quantification of ATP levels in culture wells is a convenient way to measure cytopathic effects associated to viral growth.

**Figure 1.**
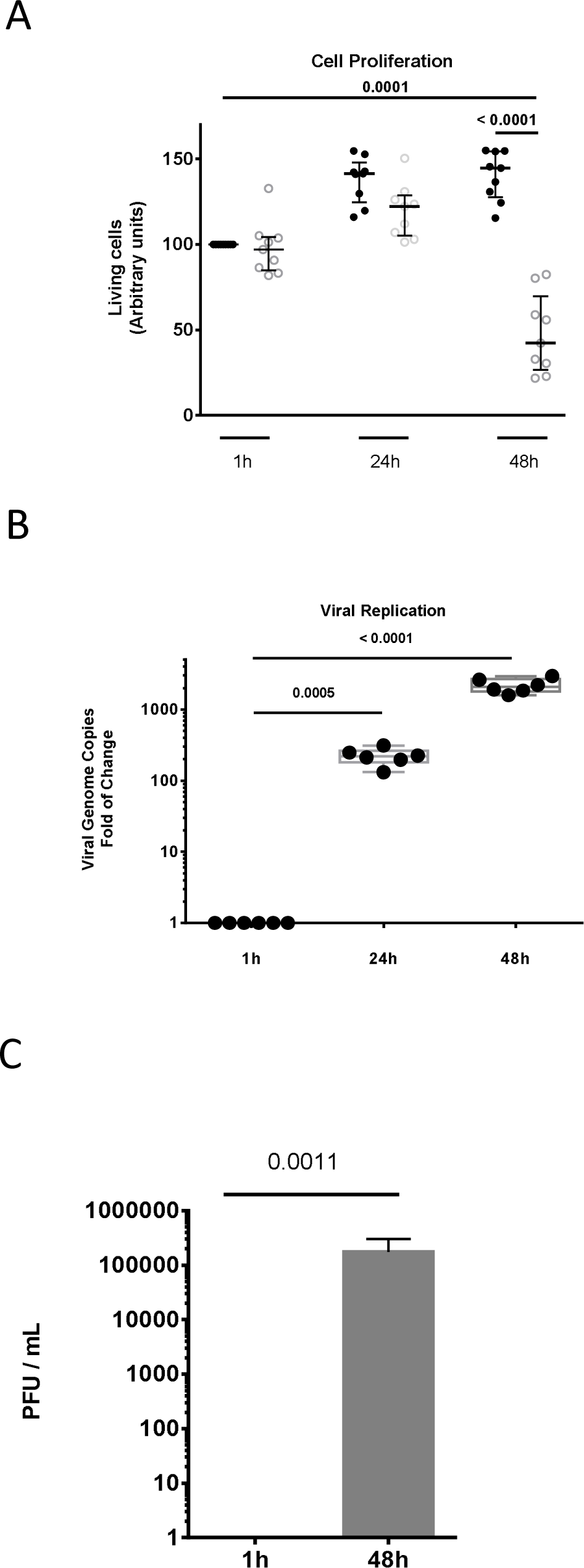
Development of a miniaturized *in vitro* infection model for EAV. (A) Equine dermal cells were mock infected (black dots) or infected with EAV at a MOI of 0.5 (white dots) and dispensed in 96-well culture plates. Cell viability was determined at 1 h, 24 h and 48 h post-infection. Each symbol represents an independent experiment (n=9). The median and interquartile range (IQR) values are shown and p values were calculate using Student’s t-test. (B) EAV genome copies were determined by one-step RT-qPCR in ED cells at 1 h, 24 h, and 48 h post-infection. Each symbol represents an independent experiment (n=6). The median and IQR values are shown. (C) Infectious particles produced by EAV-infected ED cells were quantified at 48 h post-infection on RK13 cells. Results are expressed as plaque forming units per ml of culture supernatant (PFU/ml).

### 2.2. Broad-spectrum antivirals impair EAV cytopathic effect

Previous studies on broad-spectrum antiviral molecules showed that ribavirin inhibits the replication of different nidoviruses, including *Arteriviridae* such as porcine reproductive and respiratory syndrome virus (PRRSV) ^28^. We thus tested the antiviral effects of ribarivin on EAV using the cell-viability assay described above. As a prerequisite, we first evaluated the cytotoxicity of ribavirin on ED cells. Cells were incubated with ribavirin at different concentrations (0.5, 1, 2, 5, 10, 15, 20, 50 and 100 *µ*g/ml), and the number of viable cells was determined after 1 h without treatment or after 48 h with or without treatment. No significant cell death was reported at 10 *µ*g/ml or lower concentrations of ribavirin when compared to the initial number viable cells seeded in the culture wells (Figure 2A). However, ribavirin was clearly cytostatic at these concentrations. Cell viability at 20 *µ*g/ml was 72%, and decreased to 63% and 52% when treating cells with 50 *µ*g/ml and 100 *µ*g/ml of ribavirin, respectively (Figure 2A). The half maximal toxic concentration (TC_50_), which is the concentration that kills 50% of the cells in culture, was estimated to be >100 *µ*g/ml for ribavirin. Then, we tested the effect of ribavirin in our *in vitro* EAV infection model at the non-cytotoxic concentration range of 0.5 to 20 *µ*g/ml. As shown in Figure 3A, ATP quantification in infected culture wells at 48 h showed that only 35% of cells were alive in the absence of ribavirin as opposed to 67% in presence of the drug at 0.5 *µ*g/ml. The viability of infected cells was above 80% when treated with ribavirin at 2, 5 or 10 *µ*g/ml (Figure 3A). These results were used to calculate the half maximal inhibitory concentration (IC_50_) of ribavirin that was estimated to 0.90 *µ*g/ml, *i.e.* 3.7 *µ*M. This demonstrates that ribavirin is able to protect ED cells against EAV-associated cytopathic effects.

**Figure 2.**
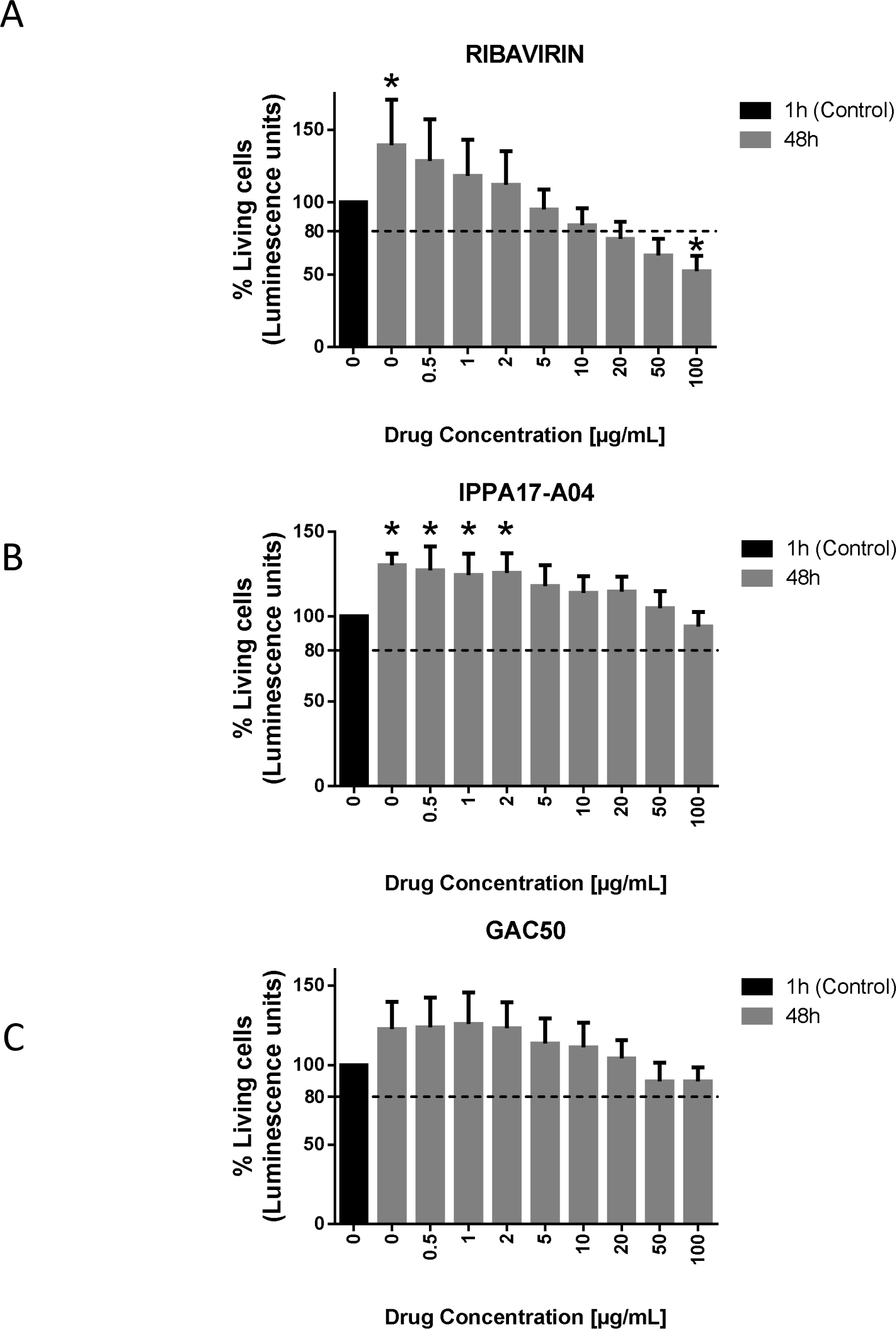
Cytotoxicity of ribavirin and the pyrimidine biosynthesis inhibitors IPPA17-A04 and GAC50. ED cells were treated with increasing concentrations of ribavirin (A), IPPA17-A04 (B) or GAC50 (C), and cellular viability was determined after 48 h of culture. The percentage of living cells relative to control wells was determined using the CellTiter-Glo reagent. Average results of four experiments are shown with standard deviation as error bars. *P value < 0.01 as calculated by one-way ANOVA with Dunnet test correction.

**Figure 3.**
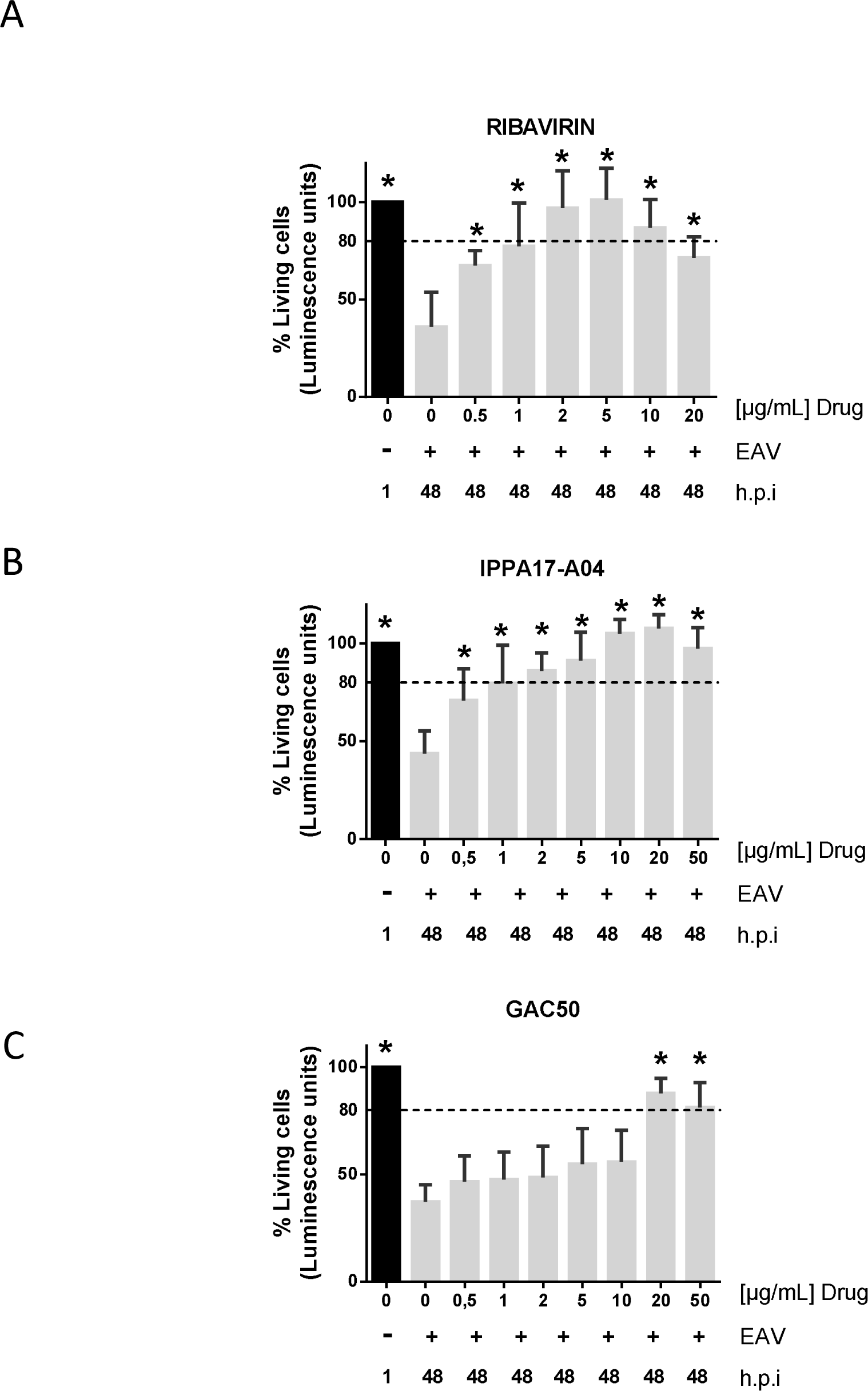
Cytopathic effects associated to EAV infection are reduced by ribavirin, IPPA17-A04 and GAC50. Cellular viability in EAV-infected cultures (MOI = 0.5) was determined when cells were treated or not with ribavirin (A), IPPA17-A04 (B) or GAC50 (C). The number of viable cells was determined after 1 h and 48 h of culture using the CellTiter-Glo reagent. Average results of four independent experiments are shown with standard deviation as error bars. *P value < 0.01 as calculated by one-way ANOVA with Dunnet test correction.

We were also interested in exploring the antiviral capacity of two new BSA, IPPA17-04 and GAC50 that impair viral replication through inhibition of *de novo* pyrimidine biosynthesis in host cells. Using our *in vitro* EAV infection assay, we tested first the cytotoxicity of both compounds. IPPA17-A04 and GAC-50 treatment did not show any sign of cytotoxicity on ED cells. At 100 *µ*g/ml, cell viability was above 80% after 48h of treatment, but cytostatic effects were observed as expected for this class of drug (Figure 2B and 2C). These results indicate that TC_50_ for IPPA17-A04 and GAC-50 is over 100 *µ*g/ml in ED cells. We then explored the cytoprotective effect of these two compounds in EAV-infected cultures at different concentrations. Our results showed that after 48 h of culture, the viability of infected ED cells reached 79% when treated with 1 *µ*g/ml of IPPA17-A04, and was above 80% when treated with concentrations >5 *µ*g/ml (Figure 3B). The IC_50_ of IPPA17-A04 was estimated at 0.70 *µ*g/ml, *i.e.* 1.74 *µ*M. GAC50 is not as effective since viability of ED infected cells treated with GAC50 is >80% only at the highest concentrations of 20 or 50 *µ*g/ml (Figure 3C) with an IC_50_ at 7.2 *µ*g/ml, *i.e.* 30.2 *µ*M. These results show that cell cultures treated with pyrimidine biosynthesis inhibitors did not exhibit cytopathic effects associated to EAV infection, suggesting that EAV replication is also impaired.

### 2.3. Ribavirin and *de novo* pyrimidine biosynthesis inhibitors impair EAV replication in ED cells

To verify that ribavirin, IPPA17-A04 and GAC50 actually block EAV replication, viral genome copies were quantified at 48 h.p.i. Culture supernatants of infected ED cells treated or not with the compounds at different concentrations were analyzed by RT-qPCR. Results showed that ribavirin at 5 and 10 *µ*g/ml significantly reduced the number of EAV genome copies in culture supernatants compared to the untreated infected cells (Figure 4A). EAV replication was totally blocked at 20 *µ*g/ml as determined by comparison with the initial inoculum quantified at 1 h.p.i. These results confirmed that ribavirin impairs EAV replication in ED cells, thus explaining the positive effect on cell survival. In parallel, we also measured viral genome copies present in the supernatant of infected cells treated with pyrimidine biosynthesis inhibitors. As expected, IPPA17-A04 and GAC50 both reduced the number of viral genome copies in culture supernatants when treated with concentrations >20 *µ*g/ml and fully blocked EAV replication at 50 *µ*g/ml (Figure 4B and 4C).

**Figure 4.**
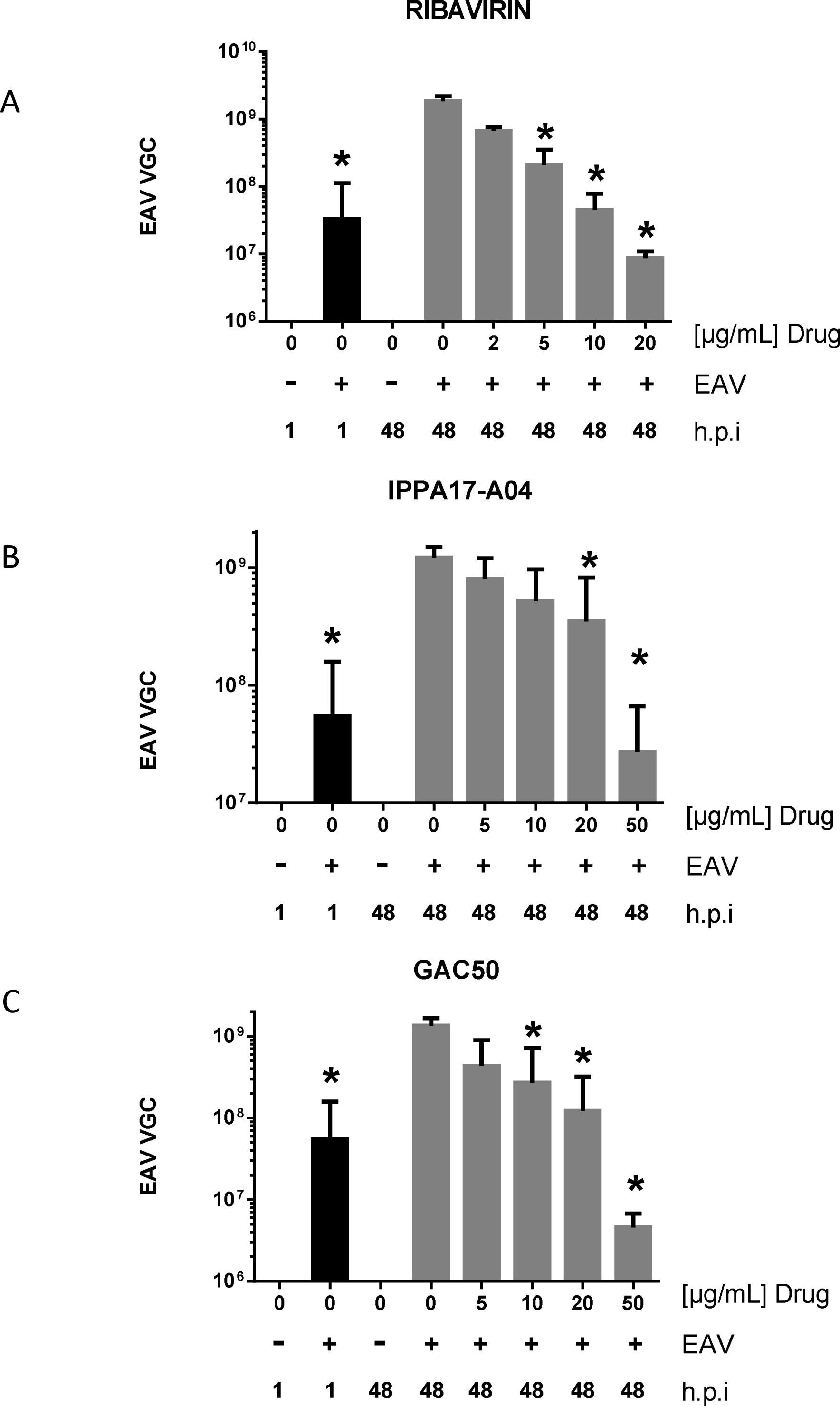
Inhibition of EAV viral replication by ribavirin, IPPA17-A04 and GAC50. Viral genome copies were quantified at 48 h post infection in supernatants of ED cells cultured in the presence or not of different concentrations of ribavirin (A), IPPA17-A04 (B) or GAC50 (C). Average results of at least 3 independent experiments are shown with standard deviation as error bars. *P value < 0.01 as calculated by one-way ANOVA with Dunnet test correction.

We checked whether compound treatments also impaired the production of newly assembled viral particles by infected ED cells. Cells were cultured in absence or presence of the compounds at different concentrations for 48 h. Supernatants were collected and viral titers determined on RK13 cells. Ribavirin treatment reduced viral titers (expressed as pfu/ml) by almost 6 logs at 5 *µ*g/ml, and more than 8 logs at 10 *µ*g/ml (Figure 5A). These results demonstrate that ribavirin sharply reduces the number of viral particles produced by infected ED cells, and thus the viral dissemination to proximal cells. On the other side, IPPA17-A04 and GAC50 showed the same antiviral activity when used at 50 *µ*g/ml with a reduction of viral titers of 7 and 8 logs, respectively (Figure 5B and 5C) whereas cytotoxic effects on ED cells were not significant (Figure 2B and 2C). This potent antiviral activity makes IPPA17-A04 and GAC50 interesting molecules to block EAV infection.

**Figure 5.**
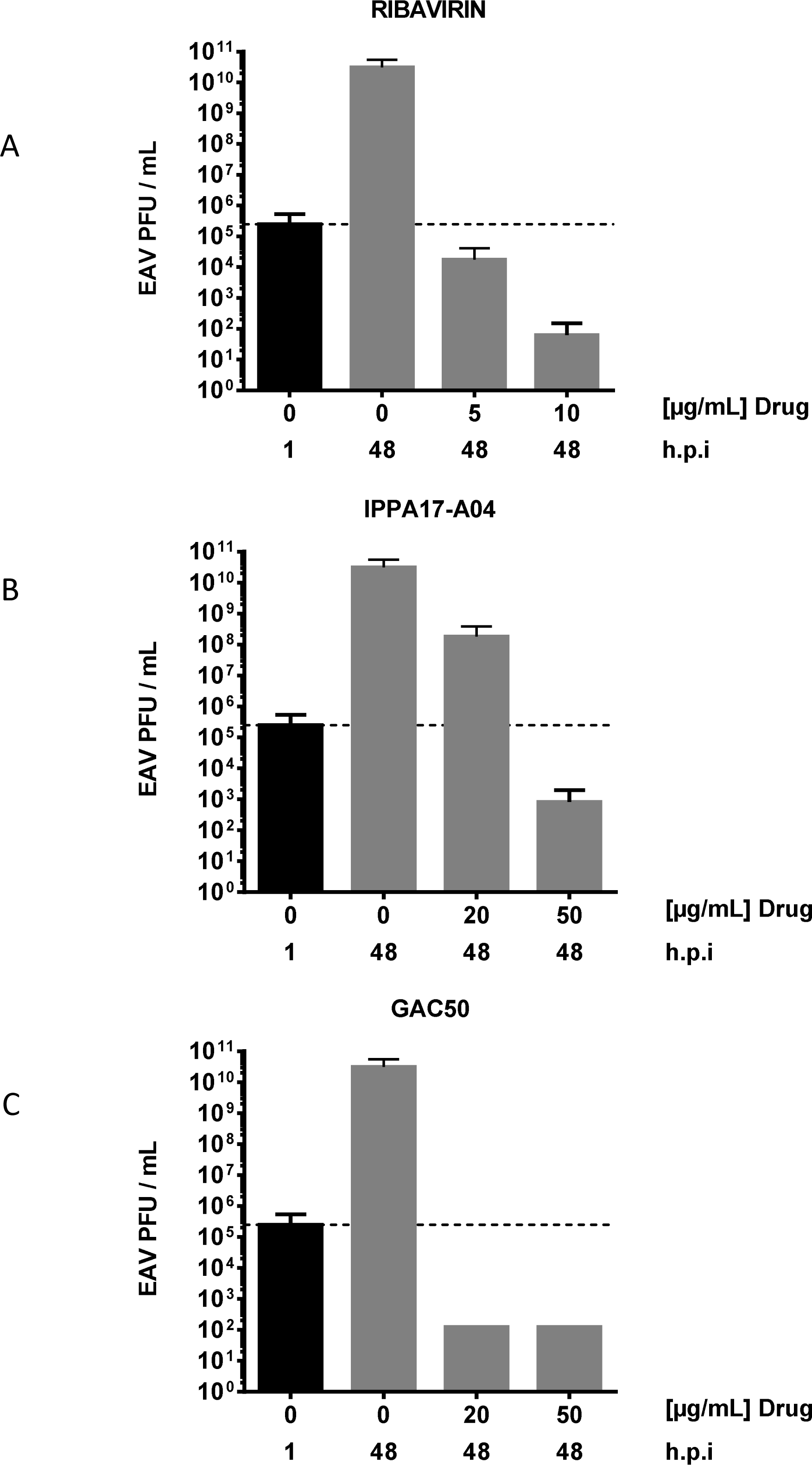
Quantification of infectious particles released by EAV-infected ED cells when treated with ribavirin, IPPA17-A04 or GAC50. Culture supernatants from EAV-infected ED cells at a MOI of 0.5 either untreated or treated with ribavirin (A), IPPA17-A04 (B) or GAC50 (C) were harvested after 48 h of culture. Infectious particles were titrated on RK13 cells and results are expressed as plaque-forming units (PFU) per ml. Average results of two independent experiments are shown with standard deviation as error bars.

### 2.4. Determination of the selectivity index for identified EAV inhibitors

To rank the three molecules that we characterized as EAV inhibitors in this study, we calculated for each molecule the Selectivity Index (SI) that corresponds to the TC_50_/IC_50_ ratio, and thus reflects the activity of a drug while taking into account cytotoxic effects. TC_50_ and IC_50_ values were determined from the dose response experiments presented in Figures 2 and 3. SI values for ribavirin, IPPA17-A04 and GAC50 were >111, >143 and >14 respectively. In conclusion, this confirms that IPPA17-A04 is, over ribavirin and GAC50, a lead molecule of interest for developing potential antiviral therapies against EAV.

### 2.5. Uridine reverses the antiviral action of IPPA17-A04

IPPA17-A04 is known to inhibit DHODH, a key enzyme involved in *de novo* pyrimidine biosynthesis. To demonstrate that IPPA17-A04 prevents EAV replication through the inhibition of *de novo* pyrimidine biosynthesis, culture medium of infected ED cells was supplemented with uridine. Indeed, uridine absorption from the microenvironment can efficiently replenish the intracellular pool of pyrimidines when *de novo* biosynthesis is impaired. Addition of uridine at 30 *µ*g/ml in the culture medium of EAV-infected ED cells abolished the cytoprotective effect of IPPA17-A04 (Figure 6A) and restored viral replication (Figure 6B). These results indicate that the antiviral activity of IPPA17-A04 in ED cells relies on the inhibition of the *de novo* pyrimidine biosynthesis pathway and depletion of the pool of pyrimidines.

**Figure 6.**
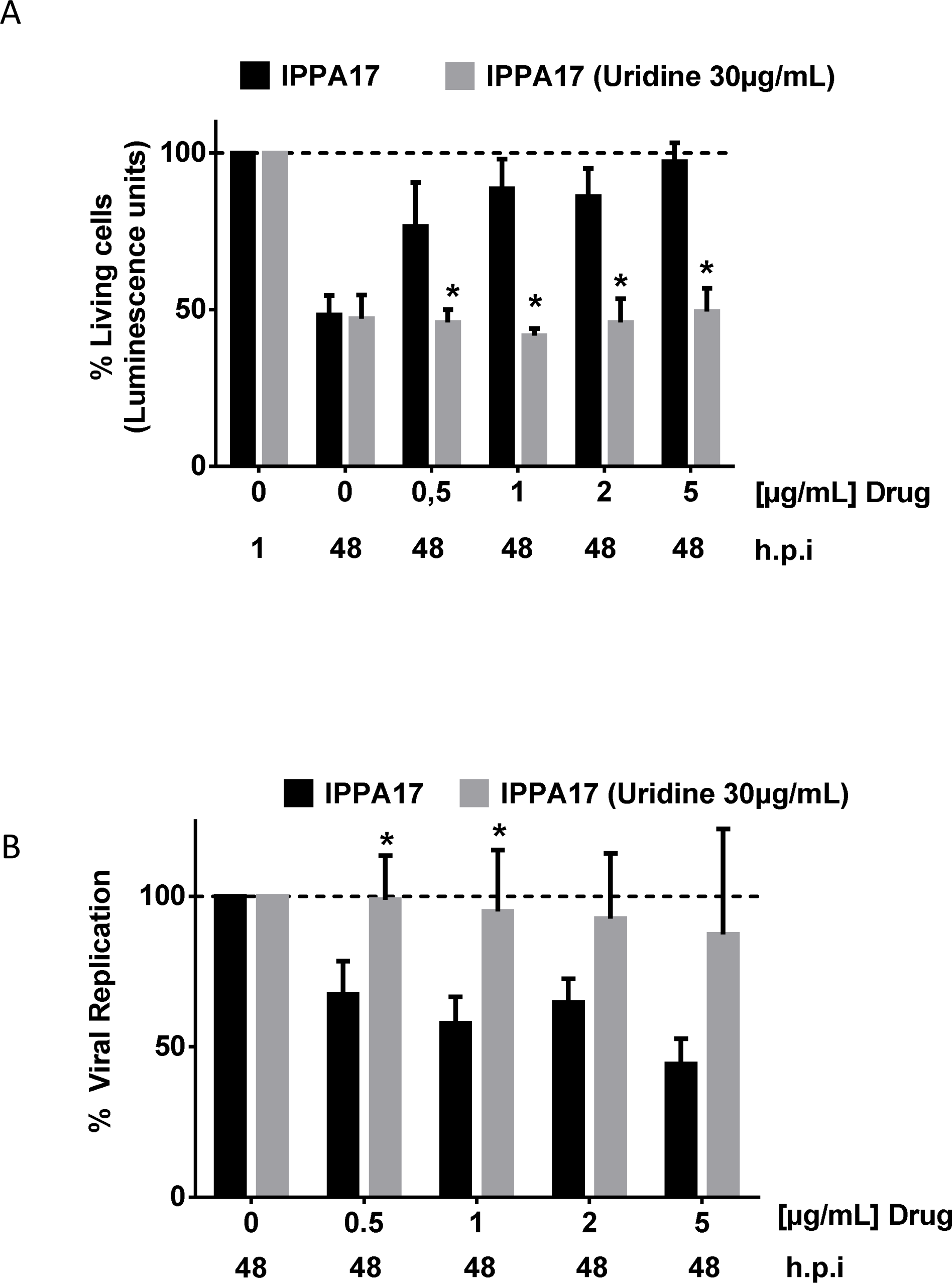
Antiviral activity of IPPA17-A04 is reversed by uridine addition in culture medium. EAV-infected cell were cultured with or without IPPA17-A04 in the absence or presence of uridine (30 *µ*g/ml). At 48 h.p.i., cell survival (A) and viral replication (B) were assessed using the CellTiter-Glo assay. Average results of three independent experiments are shown with standard deviation as error bars. * P<0.01 values were calculate using Student’s t-test.

## 3. Discussion

It remains unknown when the next EAV outbreak will occur, but since 1953, when the EAV was isolated for the first time, EAV has regularly reemerged around the world. We think that antiviral therapy used in conjunction with vaccination can reduce the viral load of infected animals, thus significantly decreasing the risk of viral dissemination in the vicinity of stables. This should also reduce the risk of animal losses due to opportunistic infections such as fulminating pneumonia in young foals. Only a recent study, exploring the inhibition of nidoviruses by the broad-spectrum antiviral molecule K22 in infected hamster cells (BHK-21), reported the first evidence of EAV inhibition by a synthetic molecule ^29^. The identification of BSA was until now impeded by the lack of suitable assay for monitoring EAV growth in high-throughput settings. To address this need, we have developed a relevant EAV infection model that is based on equine cells and was optimized for the screening of compound libraries. The infection model developed is based on the infection of the ED cell line which can be easily grown *in vitro*. This will ensure repeatability and reproducibility in HTS as opposed for example to the upper respiratory tract mucosal explant system ^30^. The goal of our study was to develop a high-throughput screening assay for evaluating the antiviral properties and the mode of action of candidate compounds against EAV.

As a proof of concept, we identified ribavirin, GAC50 and IPPA17-A04 as inhibitors of EAV in equine cells, thus paving the way for the development of therapies against EAV. IPPA17-A04 and GAC50 were not toxic for ED cells at the tested concentrations up to 100 *µ*g/ml. Among the three molecules, IPPA17-A04 was the least toxic for ED cells and the most efficient to block EAV replication as well as the production of infectious viral particles (SI>140). Addition of pyrimidine to the culture medium of EAV-infected equine cells reverted the antiviral activity of IPPA17-A04. This demonstrates that as expected for this well-characterized DHODH inhibitor, the anti-EAV effect critically depends on the inhibition of the *de novo* pyrimidine biosynthesis pathway by this molecule. The IC_50_ of IPPA17-A04 was 1.74 *µ*M against EAV in ED cells. Quite surprisingly, this is much higher than IC_50_ values previously reported for the inhibition of measles virus (MV; IC_50_ = 2.7 nM) or human respiratory syncytial virus (hRSV; IC_50_ = 5 nM) in human cell lines ^27^. This could be explained by the lower sensitivity of EAV to DHODH inhibitors. However, our data show that GAC50, another inhibitor of *de novo* pyrimidine biosynthesis, impaired EAV replication with an IC_50_ of 30.2 *µ*M, very close to hRSV in human cells (IC50 = 25 μM; ^27^). This suggests EAV is not particularly resistant to DHODH inhibitors compared to hRSV. One hypothesis to explain the lower antiviral activity of IPPA17-A04 in equine cells compared to human cells, one hypothesis is that IPPA17-A04 is less fit to equine DHODH. Indeed, despite high levels of DHODH conservation amongst mammals, and in particular between human and horse (92% of amino acid identity), host specific effects were previously reported for small molecules targeting this enzyme ^31^. For example, GAC50 but also Compound 1 described by Dong-Hoon Chung *et al* and brequinar, three unrelated DHODH inhibitor, were all shown to be active in human but not in rodent cells ^3,27,32^. This host specific profile could explain that IPPA17-A04 is more active in human cells, and well illustrates the interest of screening molecules against EAV in a culture model based on cells from equine origin. Although IPPA17-A04 already has the highest selectivity index compared to GAC50 or ribavirin, our results suggest that it could be further optimized to better fit the structure of equine DHODH and inhibit this enzyme at lower concentrations.

HTS pipelines are essential to screen large chemical libraries and/or evaluate multiple drug combinations. Here, we have developed a cost-effective, efficient, and straightforward *in vitro* EAV infection model that is fully compatible with HTS and can be automated. Because of its flexibility, our assay is appropriate for the screening of compound libraries or biological products (serum, nanobodies, cytokines, etc) and will allow the exploration of treatment combinations against EAV infection. Thanks to this assay, we demonstrated that ribavirin effectively reduces the replication of EAV in horse cells at concentrations that are not toxic. This is supports reports showing that ribavirin efficiently inhibits the replication of multiple nidoviruses from both the *Coronaviridae* and Arteriviridae family such as the MERS-CoV ^33^ and the PPRSV ^28^, respectively. Additionally, we showed that the DHODH inhibitors GAC50 and IPPA17-A04 also impair the replication of EAV, with IPPA17-A04 outperforming ribavirin and GAC50 as assessed by a higher selectivity index. It is known that Coronaviridae are inhibited by drugs targeting *de novo* pyrimidine biosynthesis, including HCoV-229E, MERS-CoV and SARS-CoV, and DHODH inhibitors were in the list of therapeutic options to be tested against SARS-CoV-2 ^34-36^. Our results demonstrate that this antiviral activity extends to *Arteriviridae*, suggesting that pyrimidine biosynthesis inhibitors may act as pan-*nidovirales* inhibitors. In conclusion, we believe that IPPA17-A04 is an interesting molecule for developing a therapy against EAV and *nidovirales* and should be further evaluated *in vivo*.

## 4. Methods

### 4.1. Compounds and dilution

Ribavirin and uridine were from Sigma-Aldrich. IPPA17-A04 and GAC50 were previously described and kindly provided by Dr. Yves Janin (Institut Pasteur, Paris, France) and Dr. Daniel Dauzonne (Institut Curie, Paris, France), respectively ^25-27^. Ribavirin was dissolved in PBS buffer and uridine in sterile water, whereas IPPA17-A04 and GAC50 were dissolved in DMSO. Compounds were tested at final concentrations ranging from 0 to 100 *µ*g/ml for cytotoxic assays and 0 to 50 *µ*g/ml for antiviral assays. Uridine was added to the culture medium of infected cells at a final concentration of 30 *µ*g/ml.

### 4.2. Cells and virus

Equine dermal cells (ED, ATTC CCL57) were used in our *in vitro* EAV infection model. ED cells were cultured at 37°C and 5% CO_2_ in DMEM (Gibco) supplemented with 1 mM of sodium pyruvate (Gibco), 50 U/ml of penicillin and 50 *µ*g/ml of streptomycin (Gibco), 10% inactivated fetal calf serum (Hyclone) and 0.025M of HEPES (Gibco). Rabbit kidney cells (RK13, ATCC CCL-37), which were used for viral titration and viral plaque assays, were cultured at 37°C and 5% CO_2_ in MEM (Gibco) supplemented with 1 mM of sodium pyruvate, 50 U/ml of penicillin and 50 *µ*g/ml of streptomycin, 1X non-essential amino acids (Gibco), 10% inactivated Fetal Calf Serum, and 0.025M of HEPES. All infection experiments were performed with a reference stock of EAV (Bucyrus strain, ATCC VR 796) titrated by plaque assay at 2.6×10^8^ PFU/ml.

### 4.3. Cell Viability Assay

Cell viability was assessed at 1, 24 and 48 h post-infection by quantification of ATP in 96-well culture plates (Greiner Bio-one) using the CellTiter-Glo Luminescent Cell Viability Assay (Promega). For each assay, displayed values correspond to the mean of 8 replicates. Luminescence was determined using a Tecan Infinite F200 PRO multimode microplate reader.

### 4.4. Infection of ED cells

ED cells were harvested using 0.25% Trypsin-EDTA (Gibco) and counted. After centrifugation, cells were resuspended in supplemented DMEM at 2.5×10^5^ cells/ml. ED cells were infected with EAV at a multiplicity of infection (MOI) of 0.5. Infected ED and mock-infected ED cells were seeded in 96-well plates (Greiner Bio-one) at 2.5.10^4^ cells per well in 100 *µ*l of supplemented DMEM and incubated for 1, 24, and 48 h. Supernatants were collected at those three-time points for viral genome quantification and virus titration.

### 4.5. RNA extraction and quantitative RT-PCR assay

RNA extraction was performed from 150 *µ*l of ED cell supernatant using the QIAamp viral RNA kit (Qiagen). EAV genome copy number was determined by one step RT-qPCR using the QuantiTect™ Virus + ROX Vial Kit (Qiagen). Each RNA sample was tested in triplicate. The primers used to quantify the EAV genome copy number are EAV ORF7F 5’-GGCGACAGCCTACAAGCTACA-3’, EAV ORF7R 5’-CGGCATCTGCAGTGAGTGA-3’ and probe is EAV ORF7P [6FAM]-TTGCGGACCCGCATCTGACCAA-[TAMRA]. The RT-qPCR program started with a retro-transcription step of 20 min at 50°C, an initial incubation step of 5 min at 95°C, and 40 cycles of a 2-step program combining 15 sec at 95°C followed by 45 sec at 60°C and fluorescence measurement. The RT-qPCR reactions were performed on a CFX Connect Real-Time PCR System (Biorad).

### 4.6. Viral titration and plaque assays

Supernatants of ED-infected cells were ten-fold serially diluted (10^−1^ to 10^−10^) in MEM (Gibco) supplemented with 1 mM of sodium pyruvate (Gibco), 50 U/ml of penicillin and 50 *µ*g/ml of streptomycin (Gibco), 0.25 M of HEPES (Gibco) and 2% inactivated Fetal Calf Serum (Hyclone). Each dilution was inoculated in duplicate on a confluent monolayer of RK13 cells seeded in 6-well culture plates. After an adsorption step of 4 h, the inoculum was removed, and 3 ml of fresh medium containing 0.75% carboxy-methyl-cellulose (Sigma-Aldrich) was added. After 3 days of culture, cell monolayers were fixed and stained with a 10% formalin buffer (Sigma-Aldrich) containing 0.1% crystal violet (Sigma-Aldrich). Presence and titration of infectious viral particles production were assessed by counting PFU.

### 4.7. Statistical analyses

Statistical analyses were performed with the GraphPad Prism software version 6.0. IC_50_ were determined using the non-linear fit « log(agonist) vs. normalized response - Variable slope».

## Acknowledgments

We would like to thank Yves Janin from the Institut Pasteur, Paris, France and Daniel Dauzonne from CNRS, UMR3666, INSERM, U1143, Institut Curie, Centre de Recherche, Paris, France who kindly provided IPPA-A14 and GAC-50, respectively. We are also very grateful to Fanny Lecouturier and Gabrielle Bouet for their technical assistance.

## Author contributions

All authors are responsible for the reported research, and all authors have participated in the conceptualization, drafting, and/or revising of the manuscript. J.C.V.C, D.G., L.M.F. and A.M. performed experiments. J.C.V.C, D.G., P.O.V. and A.H. analyzed the data and wrote the manuscript. P.D., S.P., H.M.L. and S.Z. revised the manuscript. A.H. supervised the study.

## Ethical approval

Not required

